# Cytoskeletal adaptation following long-term dysregulation of actomyosin in neuronal processes

**DOI:** 10.1101/2023.08.25.554891

**Authors:** Bruno A. Cisterna, Kristen Skruber, Makenzie L. Jane, Caleb I. Camesi, Ivan D. Nguyen, Peyton V. Warp, Joseph B. Black, Mitchell T. Butler, James E. Bear, Tracy-Ann Read, Eric A. Vitriol

## Abstract

Microtubules, intermediate filaments, and actin are cytoskeletal polymer networks found within the cell. While each has unique functions, all the cytoskeletal elements must work together for cellular mechanics to be fully operative. This is achieved through crosstalk mechanisms whereby the different networks influence each other through signaling pathways and direct interactions. Because crosstalk can be complex, it is possible for perturbations in one cytoskeletal element to affect the others in ways that are difficult to predict. Here we investigated how long-term changes to the actin cytoskeleton affect microtubules and intermediate filaments. Reducing F-actin or actomyosin contractility increased acetylated microtubules and intermediate filament expression, with the effect being significantly more pronounced in neuronal processes. Changes to microtubules were completely reversible if F-actin and myosin activity is restored. Moreover, the altered microtubules in neuronal processes resulting from F-actin depletion caused significant changes to microtubule-based transport, mimicking phenotypes that are linked to neurodegenerative disease. Thus, defects in actin dynamics cause a compensatory response in other cytoskeleton components which profoundly alters cellular function.

## Introduction

Actin is a protein that forms polarized, linear polymers that are the principal force-generating agents within a cell. One of the main functions of the actin cytoskeleton is to control cell shape by modulating cortical tension in response to vital processes such as mitosis and migration^1,2^. Actin can also produce protrusive forces by polymerizing multiple actin filaments or contractile forces through engagement with molecular motors such as non-muscle myosin II^3,4^. However, actin is just one component of a cell’s cytoskeleton, which also includes microtubules and intermediate filaments. Each of these polymer networks have their own mechanical properties, assembly/disassembly dynamics, and regulatory molecules^5^. Executing normal cellular functions requires coordination of the cytoskeletal elements through various crosstalk mechanisms^6^.

Direct cytoskeletal crosstalk may occur through crosslinking proteins that bind multiple types of filaments, such as plectin^7,8^ or tau^9,10^. Some of these coupling proteins can also act as polymerases and link the growth of different networks to each other^11,12^. Cytoskeletal crosstalk can also happen through signaling molecules that bind one type of cytoskeletal element yet regulate the dynamics of another, such as the microtubule-binding RhoA activator GEF-H1, which induces actomyosin contractility when microtubules dissasemble^13^. Finally, cytoskeletal crosstalk can also occur independently of a direct molecular connection, for example when one cytoskeletal network serves as a barrier or a channel to the growth or localization of another^14^. Because both direct and indirect forms of crosstalk may happen simultaneously, it can be difficult to determine which is predominant without precise experimental approaches that differentiate between the different mechanisms.

Here, we investigated how the loss of actomyosin affected the other cytoskeletal components. Eliminating PFN1 expression, actin filament assembly, or actomyosin contractility for extended periods of time caused an increase in the number and acetylation of microtubules, with the effect being much more pronounced in neuronal processes. Knocking out PFN1 also caused an increase in the expression of the neuron-specific intermediate filament neurofilament heavy chain. Alterations to the microtubule cytoskeleton are completely reversible following the restoration of actin and myosin activity. Moreover, the altered microtubules in neuronal processes resulting from PFN1 depletion cause significant changes to microtubule-based transport, mimicking phenotypes that have been linked to neurodegenerative disease^15,16^. Thus, defects in actin dynamics cause a compensatory response in other cytoskeleton components, with profound and far-reaching effects on cellular function.

## Results and Discussion

### Reducing F-actin increases the number and acetylation of microtubules

Recently, it was shown that the actin monomer binding protein profilin 1 (PFN1) binds tubulin and alters the polymerization of microtubules^17,18^. Clearly, PFN1-mediated microtubule regulation is important, as a PFN1 deficiency in megakaryocytes causes a Wiskott–Aldrich syndrome-like platelet defect with aberrant microtubule formations^19^ and mutations in PFN1 that cause hereditary forms of amyotrophic lateral sclerosis (ALS) prevent its binding to tubulin^18^ and hinder nerve regeneration^20^. However, the evidence is conflicting regarding whether PFN1 enhances^18,20,21^ or inhibits^17,19,22^ microtubule growth. Given PFN1’s crucial role in actin assembly^23,24^, it remains to be determined how much of its *in vivo* regulation of microtubules occurs through direct or indirect crosstalk mechanisms, which may explain the conflicting data from different model systems.

We investigated the relationship between actin and microtubules in cath.A.differentiated (CAD) cells lacking PFN1 (PFN1 KO) by immunostaining for α-tubulin and actin filaments (F-actin). Previously we established that removing PFN1 expression in these cells results in approximately 50% reduction of total F-actin^25^. PFN1 KO cells had significantly more microtubules than controls (Fig. 1A, B), which matches previous reports of PFN1-deficient cells^17,19,22^. Interestingly, there was a significant inverse correlation between microtubule and F-actin levels in PFN1 KO cells, but not in controls (Fig. 1C). This suggested an indirect crosstalk mechanism occurring after a threshold value of F-actin loss had been surpassed.

**Figure 1.**
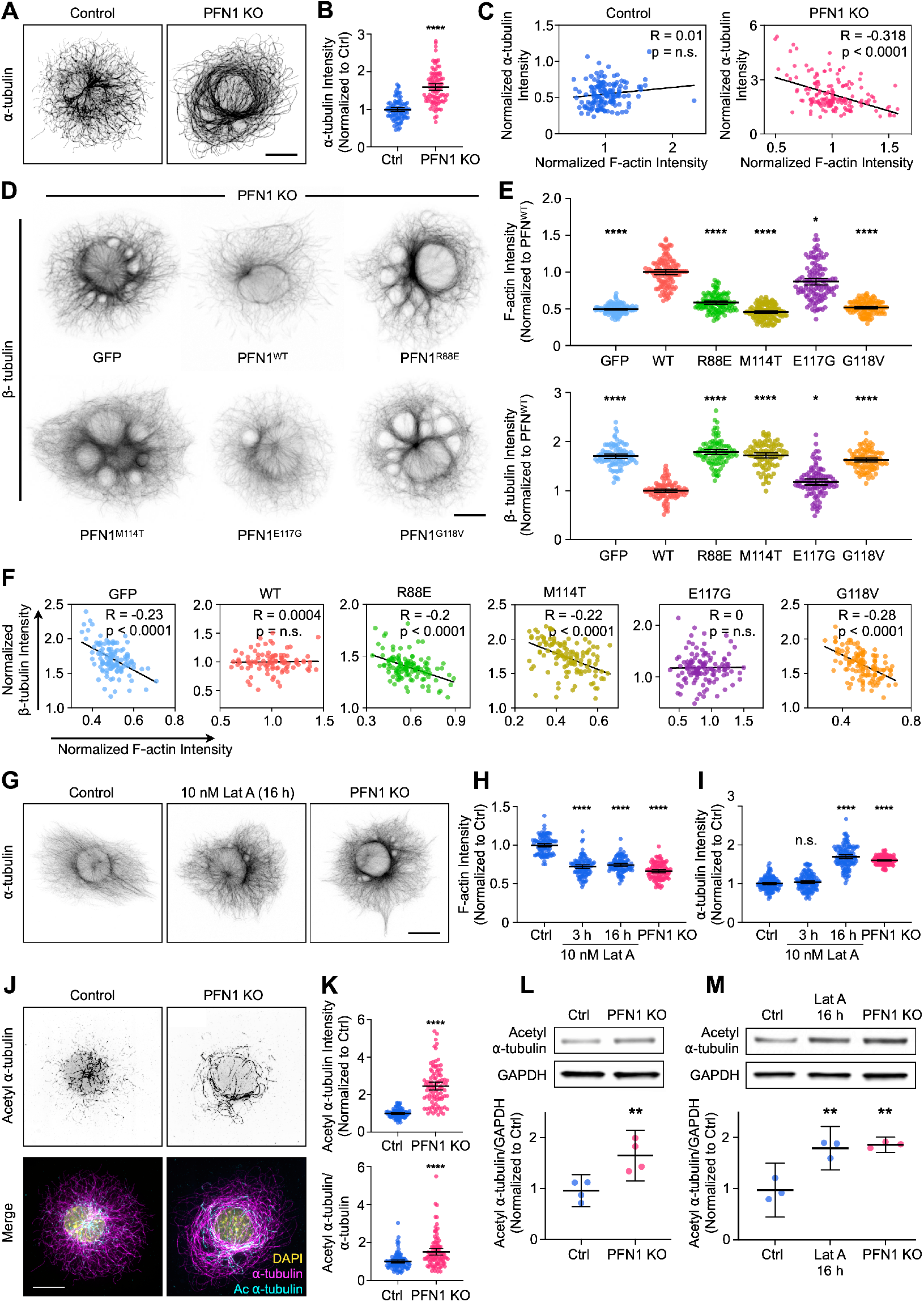
Reducing F-actin increases the number and acetylation of microtubules in undifferentiated CAD cells. **(A)** Representative images of α-tubulin in Control and PNF1 KO CAD cells. Scale bar: 10 μm. **(B)** Quantification of mean α-tubulin intensity in (A). Data are normalized to Control and plotted as mean ± 95% confidence intervals (CI). n = 97 cells for Ctrl and n = 96 cells for PFN1 KO. **(C)** Correlation between F-actin and α-tubulin intensity for cells in (A). Intensities were normalized to the mean of each dataset. **(D)** Representative images of β-tubulin in Control cells transfected with GFP, GFP-PFN1^WT^, GFP-PFN1^R88E^, GFP-PFN1^M114T^, GFP-PFN1^E117G^ or GFP-PFN1^G118V^. Scale bar: 10 μm. **(E)** Quantification of mean F-actin and β-tubulin intensity in (D). Data were normalized to PFN1^WT^ and plotted as mean ± 95% CI. n = 120 cells for each transfection. **(F)** Correlation between F-actin and β-tubulin intensity for cells in (D). Intensities were normalized to the mean of each dataset. **(G)** Representative images of α-tubulin in Control cells, Control cells incubated with 10 nM Latrunculin A (Lat A) for 16 h, and PFN1 KO CAD cells. Scale bar: 10 μm. **(H and I)** Quantification of mean fluorescence intensities in (G). F-actin (H) and α-tubulin (I). Data are normalized to Control (Ctrl) and plotted as mean ± 95% CI. n = 101 cells for each condition. **(J)** Representative images of acetyl α-tubulin in Control and PFN1 KO cells at the top and merge of DAPI, α-tubulin, and acetyl α-tubulin at the bottom. Scale bar: 10 μm. **(K)** Quantification of mean fluorescence intensities in (J). acetyl α-tubulin at the top, and the acetyl α-tubulin/α-tubulin ratio at the bottom. Data are normalized to Ctrl and plotted as mean ± 95% CI. n = 96 cells for Ctrl and PFN1 KO. **(L)** Western blot of acetyl α-tubulin in Ctrl and PFN1 KO cells at the top and quantification of levels expression at the bottom. Individual data normalized to Ctrl and plotted as mean ± 95% CI. n = 4 biological replicates for Ctrl and PFN1 KO cells. **(M)** Western blot of acetyl α-tubulin in Ctrl cells, Ctrl cells incubated with 10 nM Lat A for 16 h, and PFN1 KO CAD cells at the top, and quantification of levels expression at the bottom. Individual data normalized to Ctrl and plotted as mean ± 95% CI. n = 3 biological replicates for Ctrl and PFN1 KO cells. **** indicates p < 0.0001, ** indicates p < 0.01, * indicates p = 0.03, n.s. = not significant (p > 0.05).

To further explore whether the increase in microtubules in PFN1-deficient cells was caused by loss of direct regulation by PFN1 or indirectly through F-actin depletion^25^, we performed similar experiments in PFN1 KO cells rescued with GFP labeled wild-type (WT) and mutant PFN1, including the actin-binding deficient R88E, the ALS-causative PFN1 variants M114T and G118V, and the ALS risk factor E117G^26,27^. Previous work showed that the ALS-associated mutants, but not R88E, perturb PFN1’s ability to alter microtubule growth^18^. While WT PFN1 completely restored microtubule levels to those found in control cells, the PFN1 mutants did not (Fig. 1D, E). Interestingly, the ability of the mutants to decrease the number of microtubules was directly proportional to their ability to rescue actin polymerization (Fig. 1E). For example, E117G was able to mostly restore actin assembly and reduce microtubules, but expressing R88E, M114T, or G118V in PFN1 KO cells had essentially no effect on either parameter (Fig. 1D, E). Similarly, only WT and E117G expressing PFN1 KO cells lost the correlation between the amount of F-actin and microtubules. Interestingly, while the causative ALS-linked PFN1 mutations have been shown to alter actin assembly in complex ways, ranging from complete inhibition to enhancement over WT PFN1 in different types of polymerization assays^28,29^, their effects on total cellular levels of F-actin were remarkably similar and demonstrated a near-complete loss of function.

Since microtubule-binding proficient (R88E) and deficient (M114T and G118V) mutants failed to rescue PFN1 KO microtubule phenotypes, and restoration of microtubules was directly proportional to each PFN1 variant’s ability to rescue actin assembly, we hypothesized that the primary role of PFN1 in microtubule regulation in these cells was predominantly through actin assembly. To test this, we treated control cells with low doses of Latrunculin A (Lat A), a monomeric actin binding drug that inhibits polymerization^30^, to mimic the loss of actin filaments seen in PFN1 KO cells. We found that a treatment of 10 nM Lat A reduced actin polymerization by approximately 40% without significantly altering cell morphology (Fig. 1G). Interestingly applying Lat A to cells for three hours did not affect microtubule assembly or organization. However, an overnight Lat A treatment reproduced the PFN1 KO microtubule phenotype (Fig. 1H,I). That changes were only visible after long-term depletion of F-actin was suggestive of a homeostatic response of the microtubule cytoskeleton rather than a transient signaling mechanism induced by a microtubule-regulating F-actin binding protein.

PFN1 depletion has been shown to cause the acetylation of α-tubulin on lys40^17^. This posttranslational modification is associated with aged microtubules, and is thought to make them more curved, flexible, and resistant to breakage^31,32^. Furthermore, microtubules become hyper-acetylated in response to increased and sustained mechanical forces^33,34^, which they may be subjected to if the actin filaments that maintain cell shape and stiffness are significantly reduced^35,36^. We measured acetyl α-tubulin levels using both immunocytochemistry (Fig. 1J, K) and immunoblotting (Fig. 1L,M) and found that knocking out PFN1 or treating cells overnight with Lat A caused an identical increase in acetyl α-tubulin (Fig. 1K-M). Therefore, this change in the microtubule cytoskeleton also appears to be the result of indirect actin-microtubule crosstalk. Importantly, we measured the amount and acetylation of microtubules in control PFN1 KO mouse embryonic fibroblasts (Fig. S1) and obtained nearly identical results to those from CAD cells, indicating that these changes are not cell-type specific.

### Changes to microtubules in response to PFN1 depletion enhanced in the processes of differentiated CAD cells

Neurites are long, thin cytoskeletal structures specialized for the long-range transport of material from the cell body to the synapse. We hypothesized that the actin-microtubule homeostasis we observed in undifferentiated CAD cells would be more relevant to neuronal processes since they are principal components of this structure^37^. To test this, we first induced differentiation in CAD cells, which differentiate into a neuronal-like morphology upon serum withdrawal and form processes that are hundreds of microns long^38^. The processes of PFN1 KO cells differentiated for four days showed a similar loss of F-actin relative to controls (∼50%) (Fig. 2A, B) as to that seen in undifferentiated CAD cells (Fig. 1E). Additionally, PFN1 KO processes were narrower than controls (Fig. 2B). Despite similar decreases in actin polymerization, there was a significantly stronger increase in both the amount (Figures 2C, D, G) and acetylation of α-tubulin (Fig. 2E, F, H), in the PFN1 KO processes. While undifferentiated PFN1 KO CAD cells had an approximate 50% increase in microtubules (Fig. 1A,B) and double the amount of acetyl α-tubulin (Fig. 1J-L), PFN1 KO neuronal-like processes had a ∼400% increase in the number of microtubules (Fig. 2C, D) and triple the amount of acetyl α-tubulin as controls (Fig. 2E, F). This highlighted the importance that morphology has in the response to cytoskeletal dysregulation.

**Figure 2.**
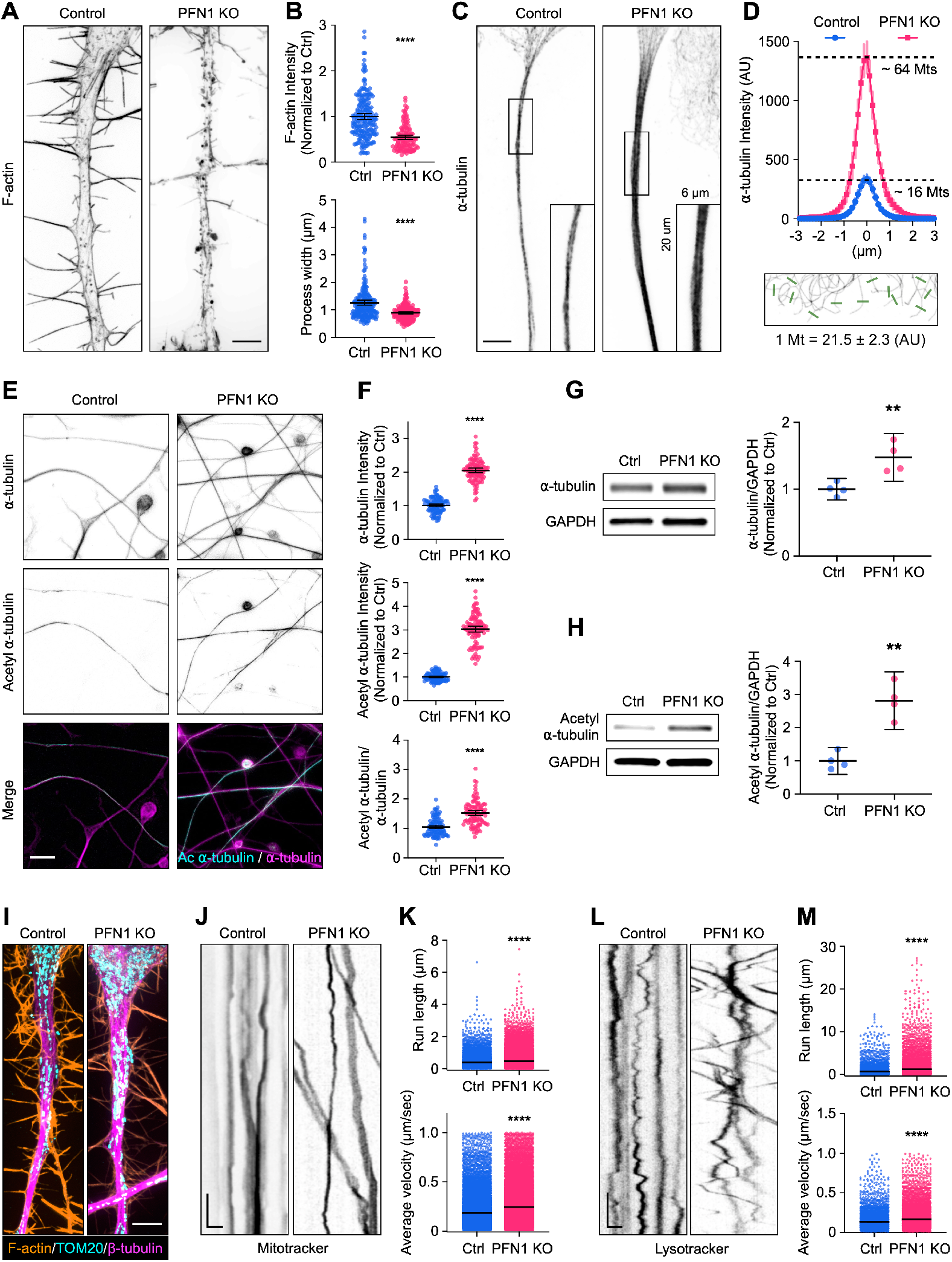
The increase in acetylated microtubules caused by PFN1 knockout is enhanced in the neuron-like processes of differentiated CAD cells and alters the active transport of organelles. **(A)** Representative images of the F-actin staining in the nascent processes of differentiated CAD cells in Control and PNF1 KO cells. Scale bar: 4 μm. **(B)** Fluorescence quantification of (A). Mean F-actin intensity at the top and Process width at the bottom. Data are normalized to Ctrl (F-actin) and plotted as mean ± 95% CI. For F-actin quantification, n = 200 processes for Control and n = 120 for PNF1 KO. For Process width quantification, n = 200 processes for Control and n = 180 for PNF1 KO. **(C)** Representative images of α-tubulin in the nascent process of differentiated CAD cells in Control and PNF1 KO cells. Scale bar: 4 μm. **(D)** Fluorescence quantification of the processes inside the rectangles in (C). The mean fluorescence intensity of ten transversal lines to the axis of each process was plotted. n = 60 processes for Control and PNF1 KO cells. The results were compared to linescans of individual microtubules from undifferentiated cells on the same coverslip (bottom image) to estimate the number of microtubules (Mts) per process. Below the image is the average fluorescence intensity of one microtubule shown in arbitrary units (AU) ±95% CI. **(E)** Representative images of the α-tubulin and acetyl α-tubulin and merge images in the processes of differentiated Control and PNF1 KO CAD cells. Scale bar: 15 μm. (**F)** Fluorescence quantification of (E). Mean α-tubulin intensity at the top, mean acetyl α-tubulin intensity in the middle, and acetyl α-tubulin/α-tubulin ratio at the bottom. Data are normalized to Ctrl (α-tubulin and acetyl α-tubulin) and plotted as mean ± 95% CI. n = 101 fields for each condition. **(G-H)** Western blot of α-tubulin (G) and acetyl α-tubulin (H) in Ctrl and PFN1 KO cells at the left and quantification of levels expression at the right. Individual data were normalized to Ctrl and plotted as mean ± 95% CI. n = 4 biological replicates for Ctrl and PFN1 KO cells. **(I)** Representative images of F-actin staining, TOM20, and β-tubulin immunostainings in processes of differentiated CAD cells in Control and PNF1 KO cells. Scale bar: 4 μm. **(J)** Kymographs from mitochondria (Mitotracker) and **(L)** lysosome (Lysotracker) in processes of differentiated CAD cells in Control and PNF1 KO cells. Vertical scale bar: 5 s and horizontal scale bar: 5 μm. **(K and M)** Kymograph quantifications. Run length at the top and Average velocity at the bottom. Data are plotted as mean ± 95% CI. For mitotracker Run length quantification, n = 7,376 for Control and n = 26,016 for PFN1 KO. For mitotracker Average velocity quantification, n = 24,441 for Control and n = 42,989 for PNF1 KO. For Lyostracker Run length quantification, n = 4,888 for Control and n = 7,534 for PFN1 KO. For Lysotracker Average velocity quantification, n = 4,937 for Control and n = 7,612 for PNF1 KO. **** indicates p < 0.0001, ** indicates p < 0.01.

An important function of neurites is to transport material from the cell body to distal regions using microtubule motors. Since microtubule acetylation can alter the binding and transport of microtubule motors^39,40^, we investigated whether changes to microtubules following long-term depletion of polymerized actin had functional consequences in the processes of differentiated PFN1 KO cells. We performed live cell imaging experiments to measure organelle transport. Despite having a similar distribution of mitochondria (Fig. 2I), there were significant differences in mitochondria mobility in the processes of PFN1 deficient cells (Fig. 2J. K), with PFN1 KO mitochondria exhibiting increased velocities and run lengths (Fig. 2K). Similar results were obtained in experiments measuring the mobility of lysosomes (Fig. 2L, M). Interestingly, organelle motility was barely affected in undifferentiated CAD cells^41^, highlighting how neuronal processes are specifically vulnerable to alterations to the microtubule cytoskeleton. Furthermore, enhanced axonal transport has been linked to neurodegenerative diseases like ALS^42,43^, Parkinson’s^44,45^ and Alzheimer’s disease^46,47^. Here we demonstrate that these processes can be significantly affected even when the microtubule cytoskeleton or its motor proteins are not directly targeted.

### PFN1 KO cells have elevated expression of neurofilament heavy chain

Neurofilaments are neuron-specific intermediate filaments that are a primary structural element of axonal processes^48^. Previous RNAseq data revealed that CAD cells express all four subunits of neurofilament (neurofilament light chain, neurofilament medium chain, neurofilament heavy chain, and peripherin)^25^. Immunocytochemistry experiments confirmed that neurofilament heavy chain was expressed in CAD cells (Fig. 3A, B). In differentiated CAD cells, it is found throughout the entire process shaft (Fig. 3A). Despite not having a significant change in *NEFH* gene expression^25^, undifferentiated PFN1 KO cells showed a two-fold increase in neurofilament heavy chain protein (Fig. 3C), with numerous fibrous accumulations in the perinuclear region (Fig. 3B). The processes of differentiated CAD cells had a similar increase in neurofilament heavy chain (Fig. 3C). Therefore, PFN1 depletion also affects the intermediate filament cytoskeleton, though unlike microtubules there was no process-specific increase. To be fair, this is not conclusive as we only investigated one neurofilament subunit and not other types of intermediate filaments. Interestingly, neurofilament accumulations are a hallmark of ALS^49,50^, which highlights another potential mechanism that PFN1 mutants could cause the disease.

**Figure 3.**
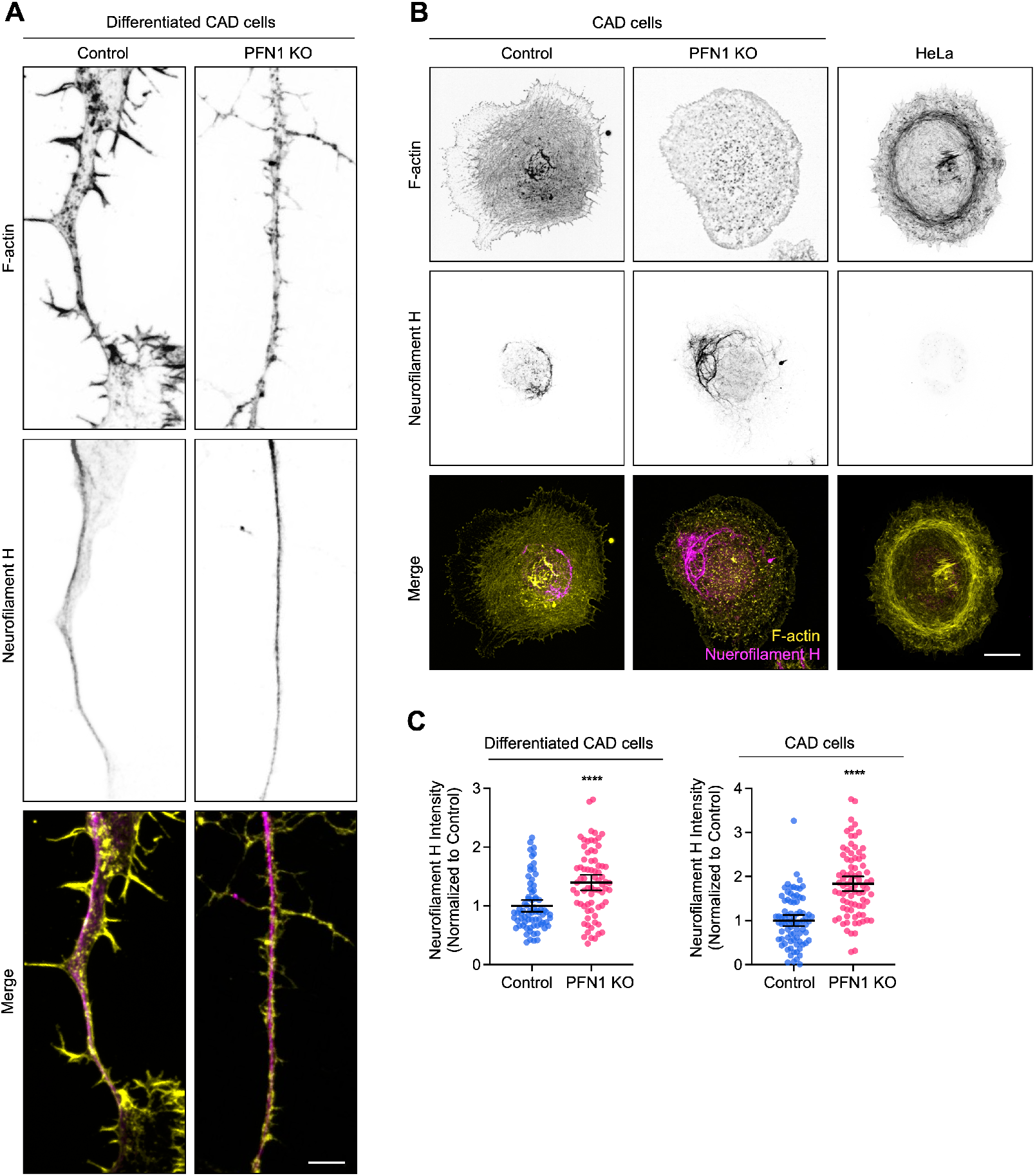
PFN1 KO cells have elevated expression of neurofilament heavy chain. **(A)** Representative images of F-actin, neurofilament heavy chain (Neurofilament H), and merge images in neuron-like processes of Control and PFN1 KO CAD cells. Scale bar: 4 *μ*m. **(B)** Representative images of F-actin, Neurofilament H, and merge images in Control, PFN1 KO CAD, and HeLa cells. Scale bar: 10 *μ*m. **(C)** Fluorescence quantification of neuron-like processes of differentiated CAD cells (A). Mean Neurofilament H intensity. Data are normalized to Control and plotted as mean ± 95% CI. n = 73 processes for Control and PFN1 KO. **(D)** Fluorescence quantification of CAD cells (B). Mean Neurofilament H intensity. Data are normalized to Control and plotted as mean ± 95% CI. n = 80 cells for Control and n = 82 cells for PFN1 KO. **** indicates p < 0.0001.

### Reducing F-actin increases the number and acetylation of microtubules in the processes of hippocampal neurons

Neurites We next wanted to determine if the same relationship between F-actin and microtubules existed in the processes of primary neurons. We cultured mouse E14 hippocampal neurons, treated them overnight with different concentrations of Lat A, and measured the relative amounts of F-actin, α-tubulin, and acetyl α-tubulin (Fig. 4A). Mimicking the results we obtained in CAD cells, we found that the number of microtubules within processes was inversely proportional to the amount of polymerized actin (Fig. 4A-C). Additionally, α-tubulin acetylation also increased as the number of actin filaments decreased (Fig. 4A, B, D), with the inverse correlation between local levels of F-actin and α-tubulin acetylation in neurites being the strongest observed in this study (Fig. 4D). While it could be argued that the processes of PFN1 KO cells have so many microtubules because they began differentiation with an increased amount of them, these experiments demonstrate that microtubules can be substantially increased in mature neurites following overnight partial actin depolymerization. Furthermore, finding a near-identical phenotype in Lat A treated hippocampal neurons addresses the concern that results obtained from CAD cells were caused by differential expression of microtubule regulators during the differentiation process.

**Figure 4.**
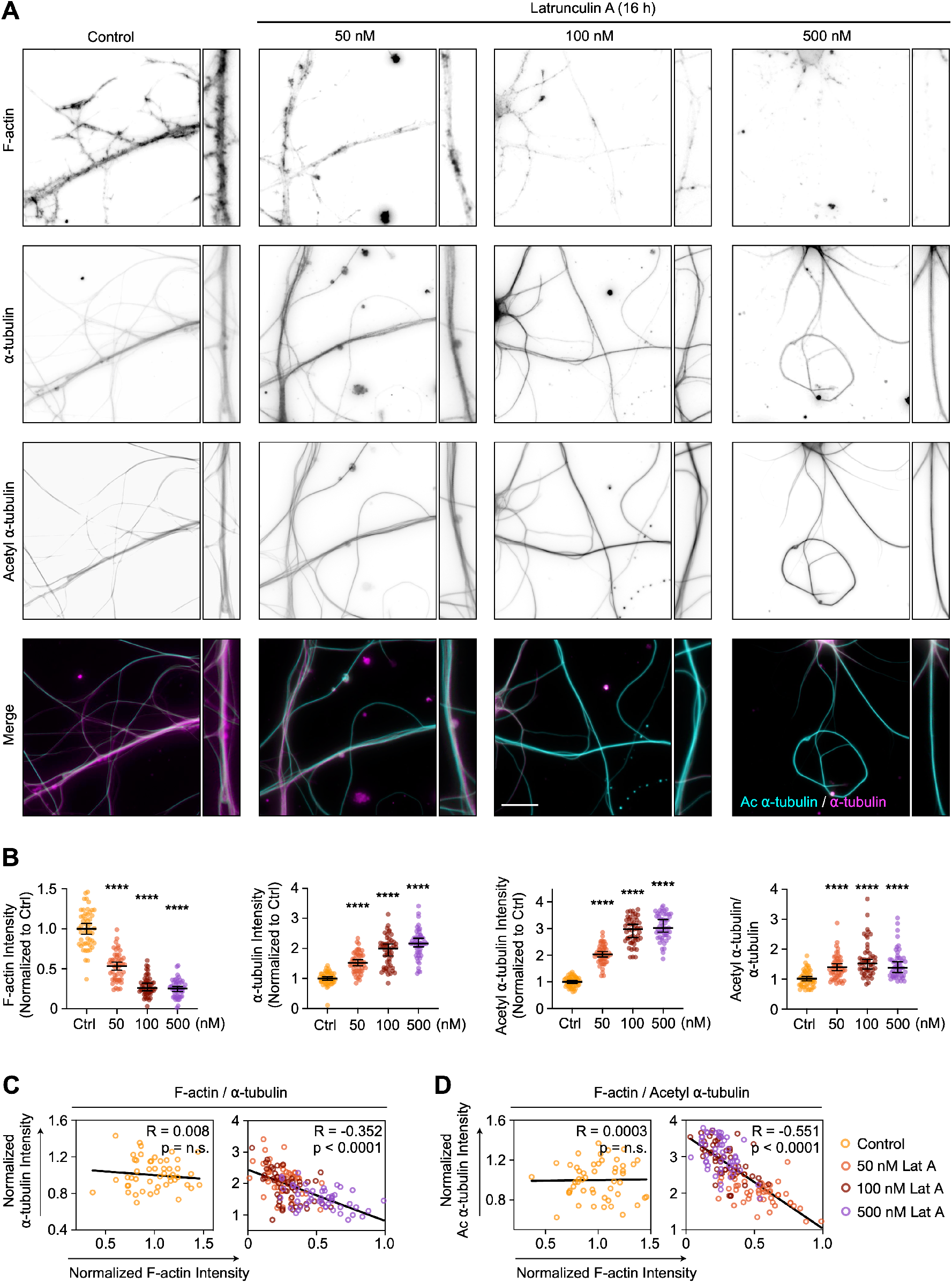
Reducing F-actin increases the number and acetylation of microtubules in the processes of hippocampal neurons. **(A)** From top to bottom, representative images of F-actin, α-tubulin, and acetyl α-tubulin, and acetyl α-tubulin/α-tubulin merge images in processes of hippocampal neurons incubated with 0, 50, 100, or 500 nM Latrunculin A (Lat A) for 16 h. Scale bar: 15 μm. **(B)** Quantification of the mean fluorescence intensity in (A). From top to bottom, F-actin, α-tubulin, acetyl α-tubulin, and acetyl α-tubulin/α-tubulin ratio. Data are normalized to Ctrl (F-actin, α-tubulin, and acetyl α-tubulin) and plotted as mean ± 95% CI. n = 50 fields for each condition. **(C and D)** Correlation between fluorescence intensities for neurites in (A). F-actin versus α-tubulin intensity (C), and F-actin versus acetyl α-tubulin (D). Intensities were normalized to the mean of each dataset. **** indicates p < 0.0001, n.s. = not significant (p > 0.05).

### Inhibition of actomyosin contractility increases the number and acetylation of microtubules without depolymerizing F-actin

Since effects on microtubules were only seen after overnight or permanent depletion of polymerized actin, we postulated that the microtubule cytoskeleton had adapted to long-term changes in the cells’ mechanical properties^2,51^. To determine if changes in cell mechanics could affect microtubules without depolymerizing actin, we treated cells overnight with the non-muscle myosin 2 inhibitor Blebbistatin, which reduces the cells’ ability to maintain cortical tension and generate contractile actin structures^52^. Treating undifferentiated CAD cells with 15 μM Blebbistatin for 24 hrs had severe effects on cell morphology, which could be reversed after removing it and replacing the medium for 24 hrs (Fig. 5A). As expected from previous work showing the effect of non-muscle myosin 2 loss of function on microtubule acetylation^53,54^, Blebbistatin treatment caused a significant increase in α-tubulin acetylation, which completely reverted to pre-treatment levels following a 24 hr wash out (Fig. 5B). In the processes of hippocampal neurons, Blebbistatin treatment also increased the number of acetylated α-tubulin without altering F-actin levels (Fig. 5C, D). As with undifferentiated cells, the effects of myosin II inhibition on microtubules were reversed after washing out Blebbistatin for 24 hrs. Also, inhibiting actomyosin contractility removed the inverse correlation between the amount or acetylation of microtubules and actin polymerization (Figure 5E). Together, these results demonstrate how neuronal processes need myosin activity and the appropriate amounts of actin filaments to retain control over microtubules and their cargo transport. Further, we have elucidated important details about a homeostasis between microtubules and actomyosin, which the cell utilizes to respond dynamically to changes in its mechanical properties.

**Figure 5.**
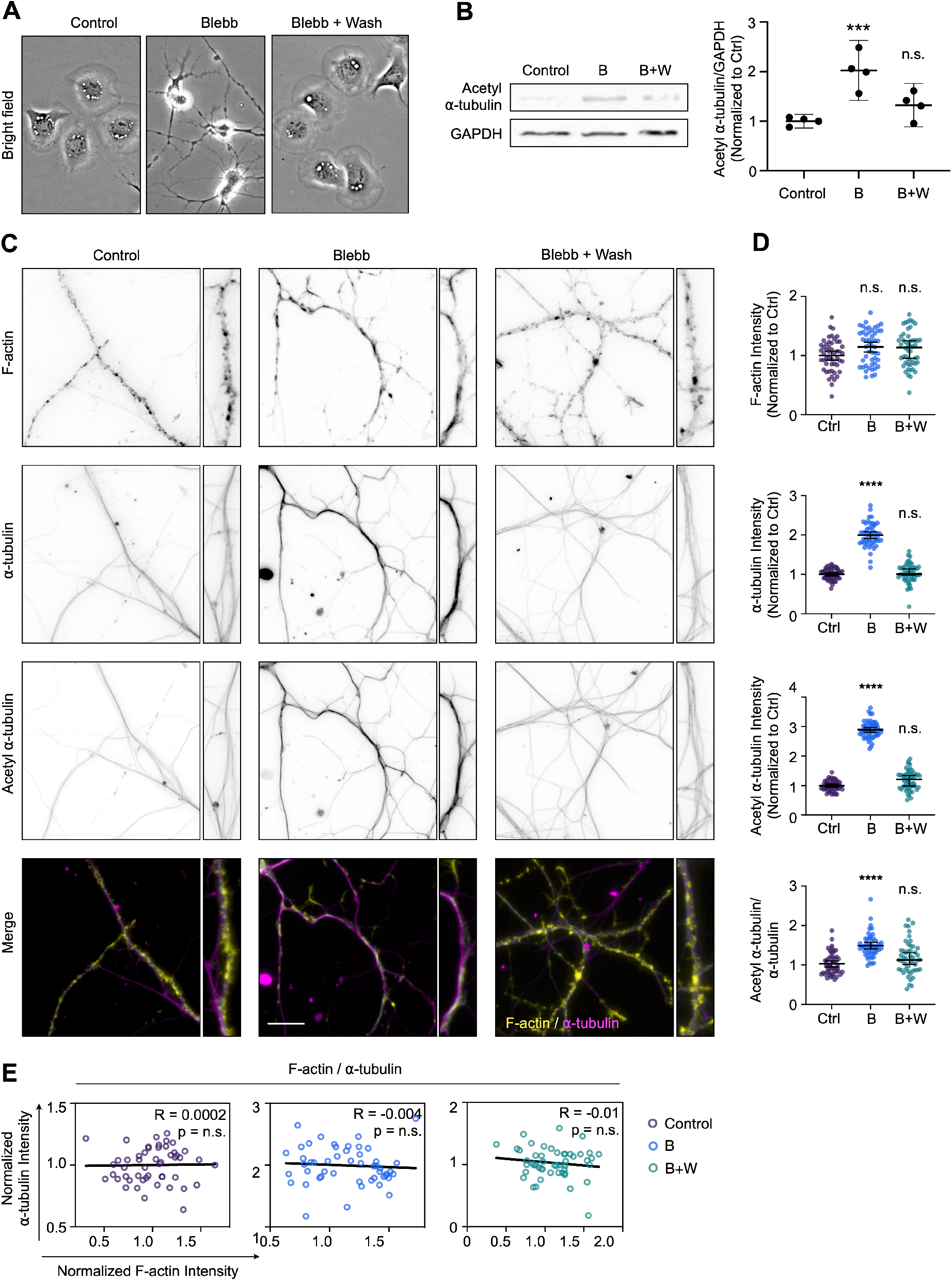
Inhibition of actomyosin contractility increases the number and acetylation of microtubules without depolymerizing F-actin. **(A)** Representative bright field images of CAD cells incubated with 0, 15 μM Blebbistatin (Blebb) for 24 h, and cells incubated with Blebb for 24 h and then washed and incubated with fresh medium (Wash) for 24 h. Scale bar: 20 μm. **(B)** Western blot of acetyl α-tubulin in CAD cells incubated with Blebb (24 h) and Blebb (24 h) plus Wash (24 h) at the top and quantification of levels expression at the bottom. Individual data normalized to Ctrl and plotted as mean ± 95% CI. n = 4 biological replicates for each condition. **(C)** From top to bottom, representative images of F-actin, α-tubulin, and acetyl α-tubulin, and F-actin/α-tubulin merge images in processes of hippocampal neurons incubated with Blebb (24 h), and Blebb (24 h) plus Wash (24 h). Scale bar: 15 μm. **(D)** Quantification of the mean fluorescence intensity in (C). From top to bottom, F-actin, α-tubulin, acetyl α-tubulin, and acetyl α-tubulin/α-tubulin ratio. Data are normalized to Ctrl (F-actin, α-tubulin, and acetyl α-tubulin) and plotted as mean ± 95% CI. n = 50 fields for Control and Blebb (24 h) plus Wash (24 h), n = 53 fields for Blebb (24 h). (E) Correlations between F-actin and α-tubulin intensity for neurites in (C). Intensities were normalized to the mean of each dataset. **** indicates p < 0.0001, n.s. = not significant (p > 0.05).

## Materials and Methods

### Cell Lines

Cath.-a-differentiated (CAD) cells: CAD cells (cat#CRL-11179, ATCC) were cultured in DMEM/F12 medium (cat#11330/032, Gibco) supplemented with 8% fetal bovine serum (FBS), 1% L-Glutamine, and 1% penicillin-streptomycin in standard tissue culture incubator. Profilin 1 knock-out (PFN1 KO) and Control (Ctrl) cells were generated from CAD cells with CRISPR/Cas9 as previously described^25^. CAD cells were differentiated under serum-free conditions for 4 days^55^.

Mouse embryonic fibroblasts (MEFs): PFN1 KO MEF clonal lines were established from previously described ARPC2 conditional knock-out mice ^56^. Among these clonal lines, JR20s were used to express Cas9 and sgRNA (5’-TCGACAGCCTTATGGCGGAC-3’) targeting mouse PFN1 ^25^ from pLentiCRISPRv2 (Addgene #52961) by lentiviral transduction. Lentivirus was generated by transfecting the plasmids pCMV-V-SVG (Addgene #8454), pRSV-REV (Addgene #12253), pMDLg/pRRE (Addgene #12251), and pLentiCRISPRv2-PFN1sgRNA (500 ng each) into HEK293FT cells using X-tremeGENE HP DNA Transfection Reagent (Sigma-Aldrich). Lentivirus was harvested at 72 hours to infect JR20 cells with 4 μg/mL of Polybrene. Around 72 hours after the lentivirus infection, JR20 cells expressing Cas9 and PFN1 sgRNA were selected using 2 μg/mL puromycin for 48 hours. LentiCRISPRv2-PFN1 (PFN1 LV) and Control cells were cultured in Dulbecco’s modified Eagle’s medium (DMEM, cat#12430-047, Gibco) supplemented with 10% FBS and 1% penicillin-streptomycin in a standard tissue culture incubator. Cells were routinely tested for mycoplasma using the LookOut Mycoplasma PCR Detection Kit (cat#MP0035, Sigma-Aldrich).

### Primary hippocampal neuron cultures

Mice primary hippocampal neuronal cultures were prepared as previously described by others ^57^. Except when otherwise indicated, all reagents were from Gibco. Briefly, hippocampi were dissected from P2 mouse pups in cold Hanks’ balanced salt solution (HBSS) supplemented with 0.08% D-glucose (Sigma-Aldrich), 0.17% Hepes, and 1% penicillin-streptomycin (Pen-Strep); filter-sterilized; and adjusted to pH 7.3. After dissection, the hippocampi were washed twice with cold HBSS and individually incubated at 37°C for 20 min in Papain dissociation solution (45 U of papain (Worthington), 0.01% deoxyribonuclease (DNase), 1 mg of DL-cysteine, 1 mg of bovine serum albumin (BSA) and 25 mg of D-glucose (all from Sigma-Aldrich) in phosphate-buffered saline (PBS). After digestion, the hippocampi were washed twice with DMEM preheated to 37°C and supplemented with 10% FBS and dissociated by 10 cycles of aspiration through a micropipette tip. Dissociated neurons were then resuspended in warm DMEM supplemented with 10% FBS and plated in 6-well plates containing 25-mm sonicated glass coverslips pretreated with 50 μg/ml poly-L-lysine (PLL, Sigma-Aldrich). After 1 hour, the medium was replaced with Neurobasal-A medium, which was supplemented with 2% B-27 and 0.25% GlutaMAX (neuronal medium). Primary neurons were maintained in a standard tissue culture incubator at 37°C with 5.5% CO_2_.

### DNA transfection

CAD cells were transfected with plasmid DNA by electroporation using the Neon Transfection System (Invitrogen) and the Neon Transfection Kit (cat#MPK1096B, Invitrogen) as previously described ^25^. Briefly, cells were grown to a confluency of 80%, trypsinized, and pelleted by centrifugation. Then, the pellet was rinsed twice with Dulbecco’s Phosphate-Buffered Saline (DPBS, cat#21-031-CV, Corning) and resuspended in a minimum amount of buffer R (Neon Transfection Kit component) with 1μg of DNA. Cells transfected with DNA constructs were cultured for 24-48 hours. Before experiments were performed, cells were grown 3 h in 10 μg/mL laminin-coated coverslips.

The following DNA constructs (Addgene) were used in this study: EGFP-C1(Plasmid #54759) and mEGFP-PFN1 (Plasmid #56438). Additional constructs used: mEGFP-PFN1-R88E (generated by us and described in Skruber et al., 2020), PFN1-ALS mutants M114T, E117G, and G118V were generated from EGFP-PFN1 plasmid with site-directed mutagenesis (Q5 New England Biolabs) using the following primers: M114T: GTC CTG CTG ACG GGC AAA GAA G (forward) and CC TTC TTT GCC CGT CAG CAG GAC (reverse), E117G: ATG GGC AAA GGA GGT GTC CAC (forward) and G GAC ACC TCC TTT GCC CAT C (reverse), G118V: ATG GGC AAA GAA GTT GTC CAC GGT GGT TTG (forward) and CAA ACC ACC GTG GAC AAC TTC TTT GCC CAT (reverse). All constructs were prepared for transfection using either the GenElute HP Endotoxin-Free Plasmid Maxiprep Kit (Sigma-Aldrich) or the NucleoBond Xtra Midi EF kit (MACHEREY-NAGEL). Correct inserts were confirmed by sequencing (Genewiz).

### Immunostaining

Cells were preheated to 37°C for 10 min, then fixed and permeabilized with 3% paraformaldehyde (Ca#15710, Electron Microscopy Sciences), 0.1% glutaraldehyde (Ca#16019, Electron Microscopy Sciences), 4% sucrose (catC12H22011, Fisher Chemical), 0.5% Triton X-100 (cat#BP151, Fisher Bioreagents), 0.1 M 1,4-Piperazinediethanesulfonic acid (PIPES, cat#4265-01, J.T. Baker) pH 7.0, 2 mM ethylene glycol-bis(2-aminoethylether)-N,N,N′,N′-tetraacetic acid (EGTA, cat#428570100, Acros Organic), 2 mM Magnesium Chloride (cat#BP214, Fisher Scientific) for 10 min at room temperature (RT). Cells were then washed three times with PBS at RT and incubated with primary antibody at 37°C for 2 hours. They were then washed four times with PBS at RT and incubated with secondary antibody for 1 h at RT, then washed four times with PBS at RT. Actin filaments were stained with Alexa Fluor 488 phalloidin,

Alexa Fluor 568 phalloidin, or Alexa Fluor 647 phalloidin (1/100 dilutions, cat#A12379; A12380; A22287; respectably, Invitrogen) in PBS for 30 min at RT. Cells were washed four times with PBS before mounting with ProLong Diamond Antifade Mountant (cat#P36961, Invitrogen). The following primary antibodies were used: rabbit anti-alpha Tubulin (1/500 dilution, cat#ab4074, Abcam), mouse anti-alpha Tubulin (acetyl K40) (1/800 dilution, cat#ab24610, clone 6-11B-1, Abcam), rabbit anti-beta Tubulin (1/500 dilution, cat#ab6046, Abcam), mouse anti-TOM20 [4F3] (1/500 dilution, cat#ab56783, Abcam). The following secondary antibodies were used: goat anti-mouse 488 (1/500 dilution, cat#A11029, Invitrogen), goat anti-rabbit 568 (1/500 dilution, cat#A11011, Invitrogen), donkey anti-rabbit 488 (1/500 dilution, cat#A32790, Invitrogen).

MEFs and CAD cells were seeded onto coverslips coated with 10 μg/mL fibronectin (cat#356008, Corning) or 10 μg/mL laminin (cat#L2020, Sigma-Aldrich), respectively, and cultured for 3 hours before fixing and permeabilization process. To perform immunostaining in hippocampal neurons, they were seeded onto coverslips coated with 50 μg/mL PLL and cultured for 3 days before the experiments.

### Imaging

Imaging was performed using the EVOS M5000 digital inverted microscope (Life Technologies) equipped with a Plan Neoflour 20X 0.5 N.A. objective and EVOS™ 40X Fluorite -LWD -0.65NA/1.79WD objective or using a Nikon CSU-W1 SoRa spinning disk confocal microscope using a 100X 1.49NA SR objective and a Hamamatsu Fusion BT Camera. All data from the SoRa had background noise removed using Denoise.ai (NIS-Elements, Nikon).

### Western Blotting

To prepare whole-cell lysates, we used a cell scraper to harvest cells in RIPA Lysis and Extraction Buffer (cat#89900, Thermo Scientific). We then passed the mixture through needles of various gauges (21, 25, and 27) five times. Next, protein quantification was assessed with Pierce BCA Protein Assay Kit (cat#23227, Thermo Scientific) and diluted in SDS buffer stained with Orange G (40% glycerol, 6% SDS, 300 mM Tris HCl, 5% β-mercaptoethanol pH 6.8). The samples were then denatured at 95°C for 5 min before loading 10 μg of samples onto a SDS-PAGE gel (Novex 4%–20% Tris-Glycine Mini Gels, cat#XP04200BOX, Thermo Scientific). Proteins were transferred to a PVDF membrane 0.2 μm (cat#10600021, Amersham) and blocked in 5% Bovine Serum Albumin (BSA, cat#A9418, Sigma-Aldrich) for 20 min. All antibodies were diluted in 5% BSA and 0.1% Tween-20 (cat#J20605AP, Fisher Scientific). The following antibodies/dilutions were used: rabbit anti-alpha Tubulin (1/500 dilution, cat#ab4074, Abcam), mouse anti-alpha Tubulin (acetyl K40) (1/800 dilution, cat#ab24610, clone 6-11B-1, Abcam), rabbit anti-GAPDH (1/1,000, cat#2118S, Cell Signaling). For secondary antibodies, goat anti-mouse (1/10,000 dilution, 2 hours at room temperature, cat#926-32210, Li-Cor) and goat anti-rabbit (1/10,000 dilution, 2 hours at room temperature, cat#926-32211, Li-Cor) were used for imaging on the Li-Cor Odyssey detection system.

### Actin depolymerization

To evaluate the effect of actin depolymerization on microtubules, we used Latrunculin A (Lat A, cat#BML-T119-0100, Enzo Life Sciences) dissolved in DMSO (cat#D2650, Sigma-Aldrich) to treat CAD cells and primary hippocampal neurons prior to immunostaining and immunoblotting. For imaging of Lat A treated CAD cells, cells were grown on laminin-coated coverslips for 3 hours, treated with 10 nM Lat A for 16 h at 37°C in a standard tissue culture incubator before immunostaining for imaging and analysis. Hippocampal primary neurons: neurons seeded onto PLL-coated coverslips and cultured for 3 days were treated with either 50 nM, 100 nM, or 500 nM Lat A for 24 h at 37°C before immunostaining for imaging and analysis. For Western blotting of CAD cells, cells cultured in 100 mm dishes to 80-90% confluence were treated with 10 nM Lat A for 16 h at 37°C before harvesting for protein electrophoresis.

### Reducing actomyosin contractility

To evaluate the role of actomyosin contractility on microtubules, we inhibited non-muscle myosin II ATPase with (-)-Blebbistatin (Blebb) (cat#20-339-11MG, MilliporeSigma Calbiochem) in CAD cells and primary hippocampal neurons. Hippocampal neurons were first seeded onto PLL-coated coverslips and cultured for 3 days, then treated with 15 μM Blebb for 24 h at 37°C, after which the medium was replaced with fresh neuronal medium for 24 h before performing immunocytochemistry, imaging, and analysis as described above. CAD cells were cultured in 100 mm dishes at 80-90% confluence, then treated with 15 μM Blebb for 24 h at 37°C after which the medium was replaced with fresh complete medium for 24 h before harvesting cells for Western blotting.

### Mitochondria and live-cell imaging

Mitotracker Red and Lysotracker green (Thermo Fisher) were used at 50 nM concentration for live cell imaging. Briefly, differentiated CAD cells grown on laminin coated coverslips for 4 days were removed from the incubator, washed once in DPBS (Gibco), and stained for 30 min at 37°C with 50nM Mitotracker or Lysotracker (dissolved in DMSO and diluted in cell culture medium without serum). After staining, cells were washed twice in a complete cell culture medium and mounted in imaging chambers with imaging media (serum- and phenol red-free DMEM/F12 with 20 mM HEPES). Mitochondria and lysosomes were imaged using total internal reflection fluorescence microscopy on a Nikon microscope using a 60X 1.49 NA Apo TIRF objective and an ORCA-Flash 4.0 sCMOS camera.

### Data analysis and statistics

Results are presented normalized to control as mean ± 95% confidence interval (CI). All data was tested for normality using the Shapiro-Wilk normality test. If the data assumed a Gaussian distribution, groups were compared using either an unpainted student’s t-test for two conditions or by ordinary one-way ANOVA followed by Tukey’s post-hoc test for comparisons of three or more conditions. If the data failed the normality test, then groups of two were compared using the Mann-Whitney test and groups of three or more were compared using the Kruskall-Wallis test followed by Dunn’s multiple comparisons test. Analysis and graphing of results were performed using Graphpad Prism software.

## Acknowledgements

Research reported in this publication was supported by the Maximizing Investigators’ Research Award from the National Institute of General Medical Sciences of the National Institutes of Health under grant number R35GM137959 to EAV.

## References

1. Chugh, P., and Paluch, E.K. (2018). The actin cortex at a glance. J Cell Sci 131. 10.1242/jcs.186254.

2. Murrell, M., Oakes, P.W., Lenz, M., and Gardel, M.L. (2015). Forcing cells into shape: the mechanics of actomyosin contractility. Nat Rev Mol Cell Biol 16, 486–498. 10.1038/nrm4012.

3. Vicente-Manzanares, M., Ma, X., Adelstein, R.S., and Horwitz, A.R. (2009). Non-muscle myosin II takes centre stage in cell adhesion and migration. Nat Rev Mol Cell Biol 10, 778–790. 10.1038/nrm2786.

4. Walker, M., Rizzuto, P., Godin, M., and Pelling, A.E. (2020). Structural and mechanical remodeling of the cytoskeleton maintains tensional homeostasis in 3D microtissues under acute dynamic stretch. Sci Rep 10, 7696. 10.1038/s41598-020-64725-7.

5. Pollard, T.D., and Goldman, R.D. (2018). Overview of the Cytoskeleton from an Evolutionary Perspective. Cold Spring Harb Perspect Biol 10. 10.1101/cshperspect.a030288.

6. Seetharaman, S., and Etienne-Manneville, S. (2020). Cytoskeletal Crosstalk in Cell Migration. Trends Cell Biol 30, 720–735. 10.1016/j.tcb.2020.06.004.

7. Fontao, L., Geerts, D., Kuikman, I., Koster, J., Kramer, D., and Sonnenberg, A. (2001). The interaction of plectin with actin: evidence for cross-linking of actin filaments by dimerization of the actin-binding domain of plectin. J Cell Sci 114, 2065–2076. 10.1242/jcs.114.11.2065.

8. Na, S., Chowdhury, F., Tay, B., Ouyang, M., Gregor, M., Wang, Y., Wiche, G., and Wang, N. (2009). Plectin contributes to mechanical properties of living cells. Am J Physiol Cell Physiol 296, C868–877. 10.1152/ajpcell.00604.2008.

9. Elie, A., Prezel, E., Guerin, C., Denarier, E., Ramirez-Rios, S., Serre, L., Andrieux, A., Fourest-Lieuvin, A., Blanchoin, L., and Arnal, I. (2015). Tau co-organizes dynamic microtubule and actin networks. Sci Rep 5, 9964 10.1038/srep09964.

10. Ramirez-Rios, S., Denarier, E., Prezel, E., Vinit, A., Stoppin-Mellet, V., Devred, F., Barbier, P., Peyrot, V., Sayas, C.L., Avila, J., et al. (2016). Tau antagonizes end-binding protein tracking at microtubule ends through a phosphorylation-dependent mechanism. Mol Biol Cell 27, 2924–2934. 10.1091/mbc.E16-01-0029.

11. Preciado Lopez, M., Huber, F., Grigoriev, I., Steinmetz, M.O., Akhmanova, A., Koenderink, G.H., and Dogterom, M. (2014). Actin-microtubule coordination at growing microtubule ends. Nat Commun 5, 4778. 10.1038/ncomms5778.

12. Henty-Ridilla, J.L., Rankova, A., Eskin, J.A., Kenny, K., and Goode, B.L. (2016). Accelerated actin filament polymerization from microtubule plus ends. Science 352, 1004–1009. 10.1126/science.aaf1709.

13. Krendel, M., Zenke, F.T., and Bokoch, G.M. (2002). Nucleotide exchange factor GEF-H1 mediates cross-talk between microtubules and the actin cytoskeleton. Nat Cell Biol 4, 294–301. 10.1038/ncb773.

14. Schaefer, A.W., Kabir, N., and Forscher, P. (2002). Filopodia and actin arcs guide the assembly and transport of two populations of microtubules with unique dynamic parameters in neuronal growth cones. J Cell Biol 158, 139–152. 10.1083/jcb.200203038.

15. Berth, S.H., and Lloyd, T.E. (2023). Disruption of axonal transport in neurodegeneration. J Clin Invest 133. 10.1172/JCI168554.

16. Millecamps, S., and Julien, J.P. (2013). Axonal transport deficits and neurodegenerative diseases. Nat Rev Neurosci 14, 161–176. 10.1038/nrn3380.

17. Nejedla, M., Sadi, S., Sulimenko, V., de Almeida, F.N., Blom, H., Draber, P., Aspenstrom, P., and Karlsson, R. (2016). Profilin connects actin assembly with microtubule dynamics. Mol Biol Cell 27, 2381–2393. 10.1091/mbc.E15-11-0799.

18. Henty-Ridilla, J.L., Juanes, M.A., and Goode, B.L. (2017). Profilin Directly Promotes Microtubule Growth through Residues Mutated in Amyotrophic Lateral Sclerosis. Curr Biol 27, 3535–3543 e3534. 10.1016/j.cub.2017.10.002.

19. Bender, M., Stritt, S., Nurden, P., van Eeuwijk, J.M., Zieger, B., Kentouche, K., Schulze, H., Morbach, H., Stegner, D., Heinze, K.G., et al. (2014). Megakaryocyte-specific Profilin1-deficiency alters microtubule stability and causes a Wiskott-Aldrich syndrome-like platelet defect. Nat Commun 5, 4746. 10.1038/ncomms5746.

20. Pinto-Costa, R., Sousa, S.C., Leite, S.C., Nogueira-Rodrigues, J., Ferreira da Silva, T., Machado, D., Marques, J., Costa, A.C., Liz, M.A., Bartolini, F., et al. (2020). Profilin 1 delivery tunes cytoskeletal dynamics toward CNS axon regeneration. J Clin Invest 130, 2024–2040. 10.1172/JCI125771.

21. Pimm, M.L., Liu, X., Tuli, F., Heritz, J., Lojko, A., and Henty-Ridilla, J.L. (2022). Visualizing molecules of functional human profilin. Elife 11. 10.7554/eLife.76485.

22. Nejedla, M., Klebanovych, A., Sulimenko, V., Sulimenko, T., Draberova, E., Draber, P., and Karlsson, R. (2021). The actin regulator profilin 1 is functionally associated with the mammalian centrosome. Life Sci Alliance 4. 10.26508/lsa.202000655.

23. Skruber, K., Read, T.A., and Vitriol, E.A. (2018). Reconsidering an active role for G-actin in cytoskeletal regulation. J Cell Sci 131. 10.1242/jcs.203760.

24. Pimm, M.L., Hotaling, J., and Henty-Ridilla, J.L. (2020). Profilin choreographs actin and microtubules in cells and cancer. Int Rev Cell Mol Biol 355, 155–204. 10.1016/bs.ircmb.2020.05.005.

25. Skruber, K., Warp, P.V., Shklyarov, R., Thomas, J.D., Swanson, M.S., Henty-Ridilla, J.L., Read, T.A., and Vitriol, E.A. (2020). Arp2/3 and Mena/VASP Require Profilin 1 for Actin Network Assembly at the Leading Edge. Curr Biol 30, 2651–2664 e2655. 10.1016/j.cub.2020.04.085.

26. Sohn, R.H., Chen, J., Koblan, K.S., Bray, P.F., and Goldschmidt-Clermont, P.J. (1995). Localization of a binding site for phosphatidylinositol 4,5-bisphosphate on human profilin. J Biol Chem 270, 21114–21120. 10.1074/jbc.270.36.21114.

27. Wu, C.H., Fallini, C., Ticozzi, N., Keagle, P.J., Sapp, P.C., Piotrowska, K., Lowe, P., Koppers, M., McKenna-Yasek, D., Baron, D.M., et al. (2012). Mutations in the profilin 1 gene cause familial amyotrophic lateral sclerosis. Nature 488, 499–503. 10.1038/nature11280.

28. Schmidt, E.J., Funes, S., McKeon, J.E., Morgan, B.R., Boopathy, S., O’Connor, L.C., Bilsel, O., Massi, F., Jegou, A., and Bosco, D.A. (2021). ALS-linked PFN1 variants exhibit loss and gain of functions in the context of formin-induced actin polymerization. Proc Natl Acad Sci U S A 118. 10.1073/pnas.2024605118.

29. Liu, X., Pimm, M.L., Haarer, B., Brawner, A.T., and Henty-Ridilla, J.L. (2022). Biochemical characterization of actin assembly mechanisms with ALS-associated profilin variants. Eur J Cell Biol 101, 151212. 10.1016/j.ejcb.2022.151212.

30. Fujiwara, I., Zweifel, M.E., Courtemanche, N., and Pollard, T.D. (2018). Latrunculin A Accelerates Actin Filament Depolymerization in Addition to Sequestering Actin Monomers. Curr Biol 28, 3183–3192 e3182. 10.1016/j.cub.2018.07.082.

31. Janke, C., and Montagnac, G. (2017). Causes and Consequences of Microtubule Acetylation. Curr Biol 27, R1287–R1292. 10.1016/j.cub.2017.10.044.

32. Eshun-Wilson, L., Zhang, R., Portran, D., Nachury, M.V., Toso, D.B., Lohr, T., Vendruscolo, M., Bonomi, M., Fraser, J.S., and Nogales, E. (2019). Effects of alpha-tubulin acetylation on microtubule structure and stability. Proc Natl Acad Sci U S A 116, 10366–10371. 10.1073/pnas.1900441116.

33. Portran, D., Schaedel, L., Xu, Z., Thery, M., and Nachury, M.V. (2017). Tubulin acetylation protects long-lived microtubules against mechanical ageing. Nat Cell Biol 19, 391–398. 10.1038/ncb3481.

34. Xu, Z., Schaedel, L., Portran, D., Aguilar, A., Gaillard, J., Marinkovich, M.P., Thery, M., and Nachury, M.V. (2017). Microtubules acquire resistance from mechanical breakage through intralumenal acetylation. Science 356, 328–332. 10.1126/science.aai8764.

35. Janke, C. (2014). The tubulin code: molecular components, readout mechanisms, and functions. J Cell Biol 206, 461–472. 10.1083/jcb.201406055.

36. Li, L., and Yang, X.J. (2015). Tubulin acetylation: responsible enzymes, biological functions and human diseases. Cell Mol Life Sci 72, 4237–4255. 10.1007/s00018-015-2000-5.

37. Coles, C.H., and Bradke, F. (2015). Coordinating neuronal actin-microtubule dynamics. Curr Biol 25, R677–691. 10.1016/j.cub.2015.06.020.

38. Kapustina, M., Read, T.A., and Vitriol, E.A. (2016). Simultaneous quantification of actin monomer and filament dynamics with modeling-assisted analysis of photoactivation. J Cell Sci 129, 4633–4643. 10.1242/jcs.194670.

39. Reed, N.A., Cai, D., Blasius, T.L., Jih, G.T., Meyhofer, E., Gaertig, J., and Verhey, K.J. (2006). Microtubule acetylation promotes kinesin-1 binding and transport. Curr Biol 16, 2166–2172. 10.1016/j.cub.2006.09.014.

40. Balabanian, L., Berger, C.L., and Hendricks, A.G. (2017). Acetylated Microtubules Are Preferentially Bundled Leading to Enhanced Kinesin-1 Motility. Biophys J 113, 1551–1560. 10.1016/j.bpj.2017.08.009.

41. Read, T.A., Cisterna, B.A., Skruber, K., Ahmadieh, S., Lindamood, H.L., Vitriol, J.A., Shi, Y., Lefebvre, A., Black, J.B., Butler, M.T., et al. (2023). The actin binding protein profilin 1 is critical for mitochondria function. bioRxiv 10.1101/2023.08.07.552354.

42. Breuer, A.C., Lynn, M.P., Atkinson, M.B., Chou, S.M., Wilbourn, A.J., Marks, K.E., Culver, J.E., and Fleegler, E.J. (1987). Fast axonal transport in amyotrophic lateral sclerosis: an intra-axonal organelle traffic analysis. Neurology 37, 738–748. 10.1212/wnl.37.5.738.

43. Hirano, A., Donnenfeld, H., Sasaki, S., and Nakano, I. (1984). Fine structural observations of neurofilamentous changes in amyotrophic lateral sclerosis. J Neuropathol Exp Neurol 43, 461–470. 10.1097/00005072-198409000-00001.

44. Prots, I., Veber, V., Brey, S., Campioni, S., Buder, K., Riek, R., Bohm, K.J., and Winner, B. (2013). alpha-Synuclein oligomers impair neuronal microtubule-kinesin interplay. J Biol Chem 288, 21742–21754. 10.1074/jbc.M113.451815.

45. Utton, M.A., Noble, W.J., Hill, J.E., Anderton, B.H., and Hanger, D.P. (2005). Molecular motors implicated in the axonal transport of tau and alpha-synuclein. J Cell Sci 118, 4645–4654. 10.1242/jcs.02558.

46. Blagov, A.V., Grechko, A.V., Nikiforov, N.G., Borisov, E.E., Sadykhov, N.K., and Orekhov, A.N. (2022). Role of Impaired Mitochondrial Dynamics Processes in the Pathogenesis of Alzheimer’s Disease. Int J Mol Sci 23 10.3390/ijms23136954.

47. Wang, Z.X., Tan, L., and Yu, J.T. (2015). Axonal transport defects in Alzheimer’s disease. Mol Neurobiol 51, 1309–1321. 10.1007/s12035-014-8810-x.

48. Yuan, A., Rao, M.V. Veeranna, and Nixon, R.A. (2017). Neurofilaments and Neurofilament Proteins in Health and Disease. Cold Spring Harb Perspect Biol 9. 10.1101/cshperspect.a018309.

49. Mizusawa, H., Matsumoto, S., Yen, S.H., Hirano, A., Rojas-Corona, R.R., and Donnenfeld, H. (1989). Focal accumulation of phosphorylated neurofilaments within anterior horn cell in familial amyotrophic lateral sclerosis. Acta Neuropathol 79, 37–43. 10.1007/BF00308955.

50. Itoh, T., Sobue, G., Ken, E., Mitsuma, T., Takahashi, A., and Trojanowski, J.Q. (1992). Phosphorylated high molecular weight neurofilament protein in the peripheral motor, sensory and sympathetic neuronal perikarya: system-dependent normal variations and changes in amyotrophic lateral sclerosis and multiple system atrophy. Acta Neuropathol 83, 240–245. 10.1007/BF00296785.

51. Li, Y., Kucera, O., Cuvelier, D., Rutkowski, D.M., Deygas, M., Rai, D., Pavlovic, T., Vicente, F.N., Piel, M., Giannone, G., et al. (2023). Compressive forces stabilize microtubules in living cells. Nat Mater 22, 913–924. 10.1038/s41563-023-01578-1.

52. Kovacs, M., Toth, J., Hetenyi, C., Malnasi-Csizmadia, A., and Sellers, J.R. (2004). Mechanism of blebbistatin inhibition of myosin II. J Biol Chem 279, 35557–35563. 10.1074/jbc.M405319200.

53. Joo, E.E., and Yamada, K.M. (2014). MYPT1 regulates contractility and microtubule acetylation to modulate integrin adhesions and matrix assembly. Nat Commun 5, 3510. 10.1038/ncomms4510.

54. Even-Ram, S., Doyle, A.D., Conti, M.A., Matsumoto, K., Adelstein, R.S., and Yamada, K.M. (2007). Myosin IIA regulates cell motility and actomyosin-microtubule crosstalk. Nat Cell Biol 9, 299–309. 10.1038/ncb1540.

55. Qi, Y., Wang, J.K., McMillian, M., and Chikaraishi, D.M. (1997). Characterization of a CNS cell line, CAD, in which morphological differentiation is initiated by serum deprivation. J Neurosci 17, 1217–1225. 10.1523/JNEUROSCI.17-04-01217.1997.

56. Rotty, J.D., Wu, C., Haynes, E.M., Suarez, C., Winkelman, J.D., Johnson, H.E., Haugh, J.M., Kovar, D.R., and Bear, J.E. (2015). Profilin-1 serves as a gatekeeper for actin assembly by Arp2/3-dependent and -independent pathways. Dev 10.1016/j.devcel.2014.10.026. Cell 32, 54–67.

57. Encalada, S.E., Szpankowski, L., Xia, C.H., and Goldstein, L.S. (2011). Stable kinesin and dynein assemblies drive the axonal transport of mammalian prion protein vesicles. Cell 144, 551–565. 10.1016/j.cell.2011.01.021.

