## Supplementary Figure 1 for "Cytoskeletal adaptation following long-term dysregulation of actomyosin in neuronal processes"

### Supplementary Figures

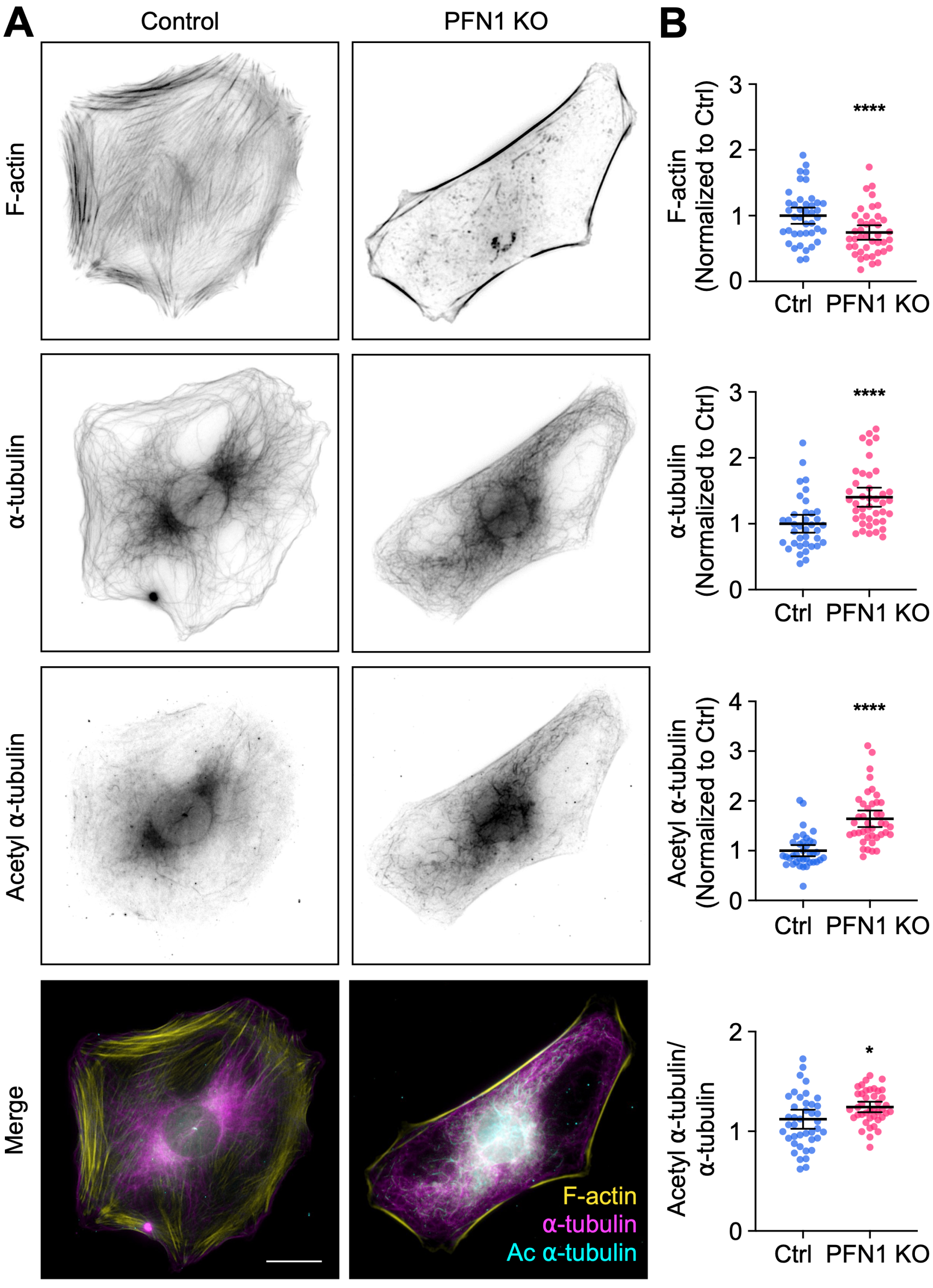


**Supplementary Figure 1. PFN1 knockout increases the number and acetylation of microtubules in mouse embryonic fibroblasts.**

**(A)** From top to bottom, representative images of F-actin staining, α-tubulin, and acetyl α-tubulin, and F-actin/α-tubulin/acetyl α-tubulin merge images in PFN1 KO mouse embryonic fibroblasts (MEFs). Scale bar: 10 µm. **(B)** Quantification of the mean fluorescence intensity in (A). From top to bottom, F-actin, α-tubulin, acetyl α-tubulin, and acetyl α-tubulin/α-tubulin ratio. Data are normalized to Ctrl (F-actin, α-tubulin, and acetyl α-tubulin) and plotted as mean ± 95% confidence intervals. n = 41 cells for each condition. **** indicates $p < 0.0001$, * indicates $p < 0.05$.
